# Brain-computer interface training fosters perceptual skills to detect errors

**DOI:** 10.1101/2025.04.26.650792

**Authors:** Deland H. Liu, Fumiaki Iwane, Minsu Zhang, Leonardo G. Cohen, José del R. Millán

## Abstract

Accurate perception of visuo-motor errors is essential for perceptual and sensorimotor learning so that corrective actions are performed timely to maintain stability and goal-directed behavior. However, a key challenge remain: conventional, behavioral perceptual training — typically based on response accuracy feedback— is limited in improving sensitivity to small, subtle errors. While prior approaches have focused on modulating sensory regions to enhance perceptual learning, we propose an alternative approach: targeting a cognitive neural marker, namely the error positivity (Pe), a component of the error-related potential (ErrP) that primarily originates in the anterior cingulate cortex (ACC), a key region involved in decision-making. We hypothesize that the Pe, which reflects conscious awareness of errors, serves as a modifiable neural correlate of error perception. To address the performance bottlenecks seen in conventional, behavioral perceptual training, we show that real-time feedback on ErrP presence or absence during perceptual training over five longitudinal days enhances the Pe component and improves participants’ ability to detect small visuo-motor errors —an outcome not achieved through conventional, behavioral perceptual training. These findings offer new neurophysiological insights into error perception and learning, and establish ErrP-based BCI interventions as a promising tool for accelerating perceptual learning, particularly in contexts where subtle error detection is critical.

---

Perceptual skills, the ability to make decisions based on ambiguous sensory information, play a critical role in detecting visuo-motor errors. These errors can arise from mismatches between intended motor commands and sensory feedback or unexpected environmental changes^1^. Accurate perception of visuo-motor errors is essential for maintaining motor accuracy and enabling adaptive responses^2–5^. For instance, perceiving small disturbances facilitates real-time corrective actions, preventing accidents or falls^6^. Such error detection can also drive sensorimotor learning, allowing individuals to refine motor skills and adapt to dynamic environments^3, 7, 8^. For instance, in split-belt walking studies, participants adapted their step length and timing after receiving inaccurate foot placement feedback, retaining these adaptations even after the perturbation was removed^9^.

Perceptual skills are adaptable and can be refined through training, a process known as perceptual learning^10, 11^. Practice-induced perceptual improvements have been reported across different sensory domains, including visual, olfaction and audition^12–16^. However, prior findings indicate that perceptual learning follows an asymptotic trajectory, eventually plateauing where further refinements become increasingly difficult^10, 17^. This suggests that while conventional, behavioral perceptual training —typically characterized by repetitive exposure and explicit behavioral feedback— may enhance the detection of large visuo-motor errors, it may be insufficient for improving sensitivity to smaller errors. Since small visuo-motor errors are crucial in precision-dependent tasks, alternative training strategies are needed to overcome perceptual constraints, as even minor deviations can have significant consequences. For instance, during surgical procedures, expert surgeons must detect and compensate for subtle visuo-motor perturbations to maintain accurate motor control, both in direct manual operations and in telerobotic surgical environments, where visual delays can exacerbate such errors ^18, 19^. In high-performance sports such as baseball, athletes rely on superior visuo-motor skills to perceive subtle variations in ball trajectory, speed, and spin and adjust their motor responses accordingly^20^. Similarly, in everyday activities, fine motor tasks such as handwriting require strong visuo-motor sensitivity^21^.

To enhance perceptual learning of small visuo-motor errors we build on the neural mechanisms underlying error detection and design an innovative strategy that leverages these mechanisms to improve learning outcomes. Critically, our approach does not target enhancement in early sensory neural visual representations, but higher decision-making processes. In the context of error perception, one electroencephalogram (EEG) event-related potential (ERP) of particular interest is the error-related potential (ErrP), which is elicited when humans perceive an erroneous action either by themselves^22, 23^, another person, or an autonomous device^24, 25^. The ErrP is characterised by two distinct components following an incorrect action: the error-related negativity (ERN), an early negative deflection, and the error positivity (Pe), a subsequent positive deflection, typically observed in the frontal, central and centro-parietal regions^5, 25–28^. While the ERN is linked to early error detection^29^, the Pe component is associated with conscious error awareness. The Pe amplitude has been shown to be higher during consciously perceived errors compared to those that are partially or not consciously recognized^26, 30^. Additionally, Pe is believed to encode the strength of internal decision evidence indicating an error has occurred^31^ and is sensitive to the magnitude of errors and expectation mismatches^32, 33^.

Building on these prior evidence, we first hypothesize that the Pe component of the ErrP is a neural correlate of error perception and can be modulated through perceptual training. To test this, we designed a visuo-motor error perception task in which participants controlled a cursor along a straight trajectory using a joystick (Figure 1A). This task is adapted from established cursor-reaching paradigms in the literature that employ visuo-motor rotations applied to continuous trajectories to study ErrPs^2, 5^. In half of the trials, varying magnitudes of visuo-motor errors (3^◦^, 6^◦^, 9^◦^, or 12^◦^) were introduced to control the difficulty levels of perception, requiring participants to make corrective actions to maintain a straight trajectory. Participants completed 5 consecutive days of perceptual training on this task. In Experiment 1, participants underwent conventional perceptual training and received behavioral feedback on their accuracy in detecting the errors. We hypothesize that conventional, behavioral perceptual training will improve perception of larger visuo-motor errors (9^◦^ and 12^◦^), but will be less effective for smaller errors (3^◦^ and 6^◦^).

**Figure 1:**
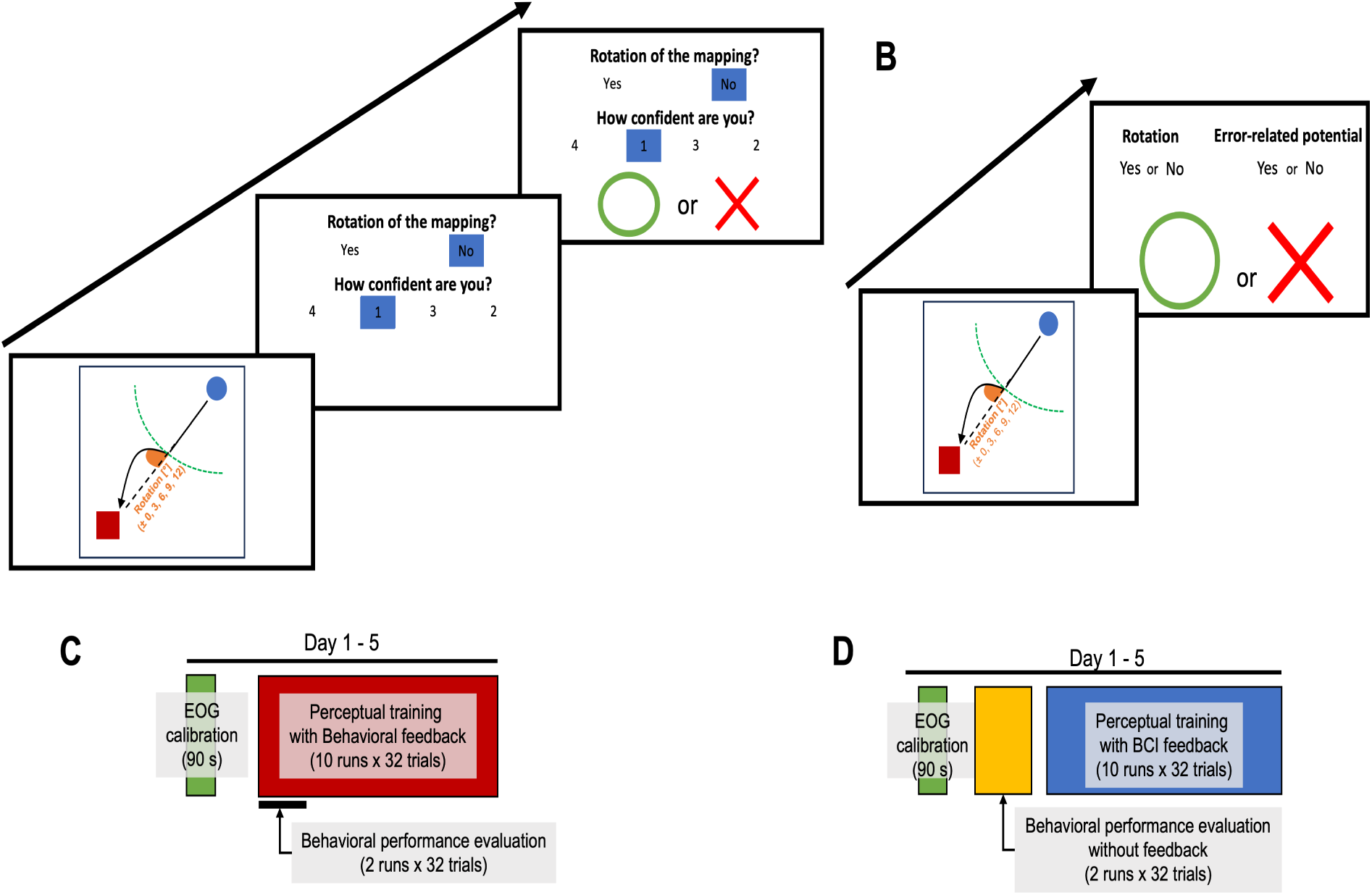
Experimental design. **A**: Participants in the Behavior group performed a cursor-reaching task using the left joystick on a gamepad, controlled by their left thumb. They were instructed to create a straight cursor trajectory for each trial, which could be perturbed by a visuo-motor rotation. The solid black line represents an example cursor trajectory, whereas the dashed black line shows an ideal straight cursor trajectory. The dashed green curve marks the invisible boundary where the normal joystick-to-cursor mapping could be violated by rotations of varying magnitudes (±0^◦^, 3^◦^, 6^◦^, 9^◦^, 12^◦^). After each trial, participants indicated whether they perceived a rotation by pressing joystick buttons (blue squares) and received behavioral feedback on their response. **B**: Participants in the BCI group completed the same task, but their feedback was the output of their BCI (detection of the presence/absence of an ErrP during the cursor trajectory) together with information about an eventual rotation. **C**: Participants in the Behavior group underwent 5 days of training, each day consisted of 90 s of EOG calibration followed by 320 trials of perceptual training (10 runs of 32 trials each). **D**: Participants in the BCI group also completed 5 days of training, identical in the number of training trials to the Behavior group, with an additional behavioral assessment of 64 trials (2 runs of 32 trials) conducted after EOG calibration and before the main training block of each day. The BCI group also performed a behavioral assessment block the day after finishing the 5-day training.

Additionally, we postulate that for errors that participants fail to learn to perceive (3^◦^ and 6^◦^) the Pe amplitudes will be lower in missed trials compared to those in successfully detected trials. To enhance the perception of small visuo-motor errors, we developed a novel brain-computer interface (BCI) strategy that provides real-time feedback on the presence or absence of ErrPs during perceptual training. We hypothesize that this approach will enhance the Pe component over the course of training and improve perception of small visuo-motor errors. To test this hypothesis, Experiment 2 employed the same visuo-motor error perception task with a different group of participants who received real-time feedback from a personalized ErrP-based BCI. This feedback included information on the presence or absence of an ErrP and the nature of the trial (correct —i.e., no rotation— or erroneous —i.e., rotation) (Figure 1B).

## Results

### Pe component reflects error magnitude and conscious error awareness of small visuo-motor errors

Figure 2, the grand-averaged event-related potential at Cz, computed across all participants from both groups, all training days, and at different rotation conditions shows that the Pe component encodes the magnitudes of errors. Pe amplitude increased progressively with the error magnitude (Behavior group: Spearman correlation, *r*(75) = 0.8457, *p <* 0.0001; BCI group: Spearman correlation, *r*(74) = 0.6820*, p <* 0.0001).

**Figure 2:**
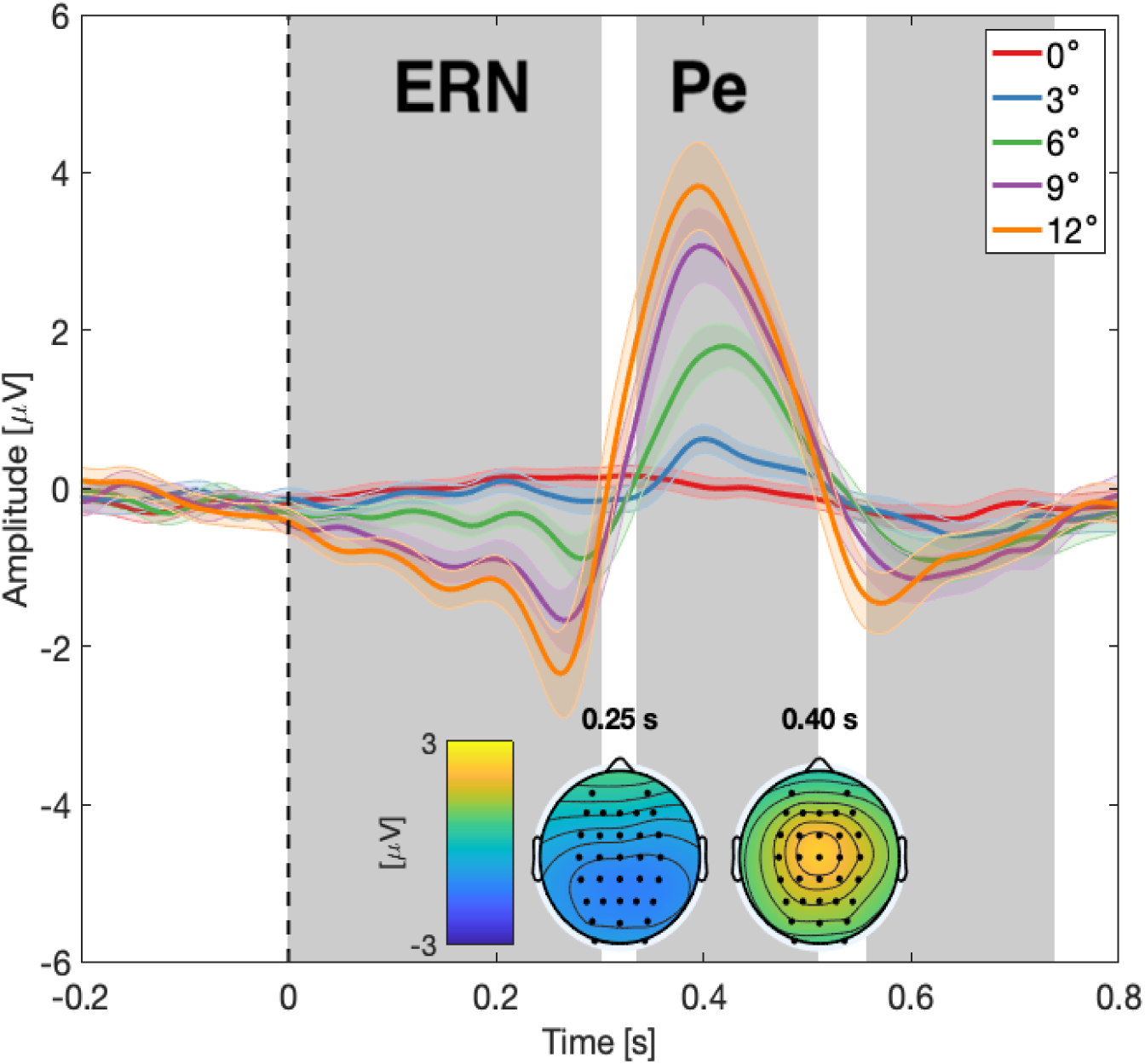
Grand average event-related potentials. Time-locked grand average EEG potentials at channel Cz of all participants in the two groups (N=32) at different rotation magnitudes (0^◦^, 3^◦^, 6^◦^, 9^◦^, 12^◦^). Time 0 s, marked by horizontal black dashed line, corresponds to the onset of rotation. The shaded areas around the EEG potentials represent the standard error over participants. Gray shaded regions highlight time intervals where significant differences were found between the error and correct class epochs. The first region, spanning approx. time window of [0, 300] ms w.r.t trigger onset, is the ERN region. The second region, spanning approx. time window of [340, 520] ms w.r.t trigger onset, is the Pe region. Insets represent topography of EEG amplitude for erroneous epochs (grand average across all subjects (N=32) and rotation magnitudes) at two different time points with respect to the rotation onset; i.e., 250 and 400 ms

To further explore how the Pe component is modulated by conscious versus unconscious error awareness, Figures 3A,B show the grand averaged signals for 3^◦^ and 6^◦^ rotations, comparing successfully perceived trials to missed trials within the Behavior group. A clear amplitude difference in the Pe region, indicated by the gray shaded area, was observed between these conditions. A two-way rmANOVA, with factors of rotation magnitude (3^◦^, 6^◦^) and error perception (successful vs. missed trials), revealed a significant main effect of error perception on Pe amplitude (*F* (1, 15) = 27.9740, *p* = 0.0001).

**Figure 3:**
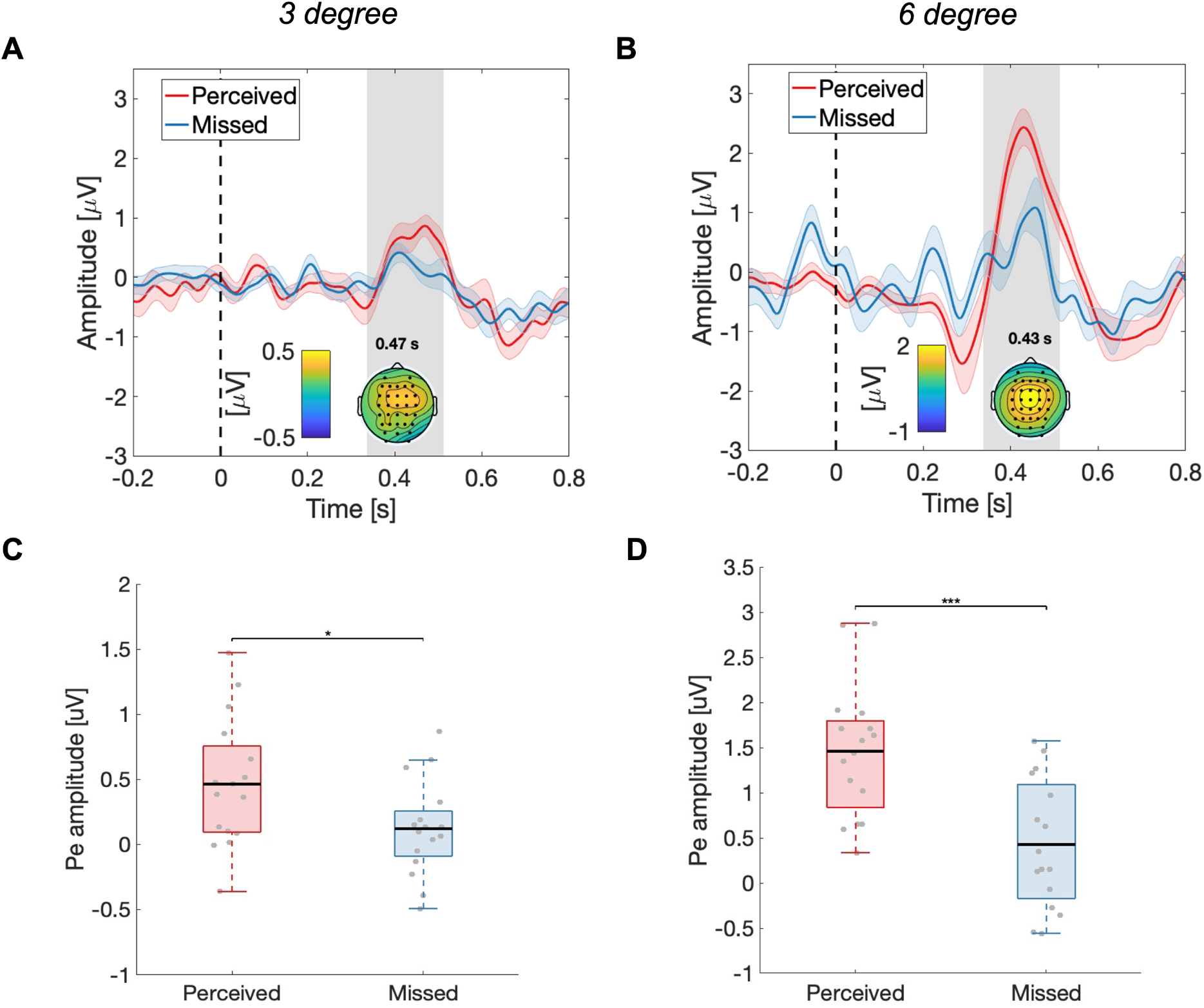
ErrPs for small visuo-motor errors: comparison of successfully perceived vs. missed trials. *A, B*: Grand-averaged ErrPs at Cz for 3^◦^ and 6^◦^ errors in the Behavior group (N=16), shown for successfully perceived trials (red) and the missed conditions (blue). The horizontal dashed black line indicates the onset of rotation. The shaded areas around the EEG potentials represent the standard error across participants. Gray shaded regions mark the Pe region. Insets display EEG amplitude topographies for the two rotations at Pe time points. *C, D*: The Pe amplitude at Cz for 3^◦^ and 6^◦^ errors in successfully perceived trials (red) and missed conditions (blue). Each boxplot represents the distribution of the Pe amplitude across participants, with gray dots representing individual participants’ mean Pe amplitudes. Pe comparisons used paired comparisons. ∗*P <* 0.05, ∗ ∗ ∗*P <* 0.001

Figures 3C, D quantify these differences using paired comparisons. For 3^◦^ rotations, the Pe amplitude was significantly higher in successfully perceived trials compared to missed trials and an effect size close to large (Successfully perceived: 0.4651 ± 0.4925 *uV*, Missed: 0.1212 ± 0.3628 *uV, t*(15) = 2.8652*, p_corrected_* = 0.0118*, Cohen*^t^*s d_z_* = 0.7950). For 6^◦^ rotations, the difference was even more pronounced with a large effect size (Successfully perceived: 1.4605 ± 0.7378 *uV*, Missed: 0.4262 ± 0.7143 *uV, t*(15) = 4.5623*, p_corrected_* = 0.0008*, Cohen*^t^*s d_z_* = 1.4245).

### Pe amplified through perceptual training

The trend in Pe amplitude across days in the Behavior group (Experiment 1), as shown in Figure 4, demonstrates that the Pe component can be modulated by perceptual training. A two-way rmANOVA revealed a significant main effect of rotation magnitude (*F* (4, 60) = 53.6290, *p <* 0.0001) and training days (*F* (4, 60) = 4.8195, *p* = 0.0019), and a significant interaction between the two (*F* (16, 240) = 2.1260, *p* = 0.0080). Linear mixed-effects analysis of Pe changes across days at each rotation magnitude showed significant enhancements for 6^◦^ (*β*(78) = 0.1508 ± 0.0515, *F* (1, 78) = 8.5578, *p_corrected_* = 0.0113, *R*^2^(80) = 0.4571), 9^◦^ (*β*(78) = 0.1656 ± 0.0656, *F* (1, 78) = 6.3715, *p_corrected_* = 0.0227, *R*^2^(80) = 0.6601), and 12^◦^ (*β*(78) = 0.3020 ± 0.0723, *F* (1, 78) = 17.4390, *p_corrected_* = 0.0004, *R*^2^(80) = 0.7752). Results at 0^◦^ approached significance (*β*(78) = 0.0467 ± 0.0240 *F* (1, 78) = 3.7811, *p_corrected_* = 0.0692, *R*^2^(80) = 0.6495), although the coefficient was near zero. At 3^◦^, no significant effect was observed (*β*(78) = 0.0426 ± 0.0387, *F* (1, 78) = 1.2075, *p_corrected_* = 0.2752, *R*^2^(80) = 0.1681).

**Figure 4:**
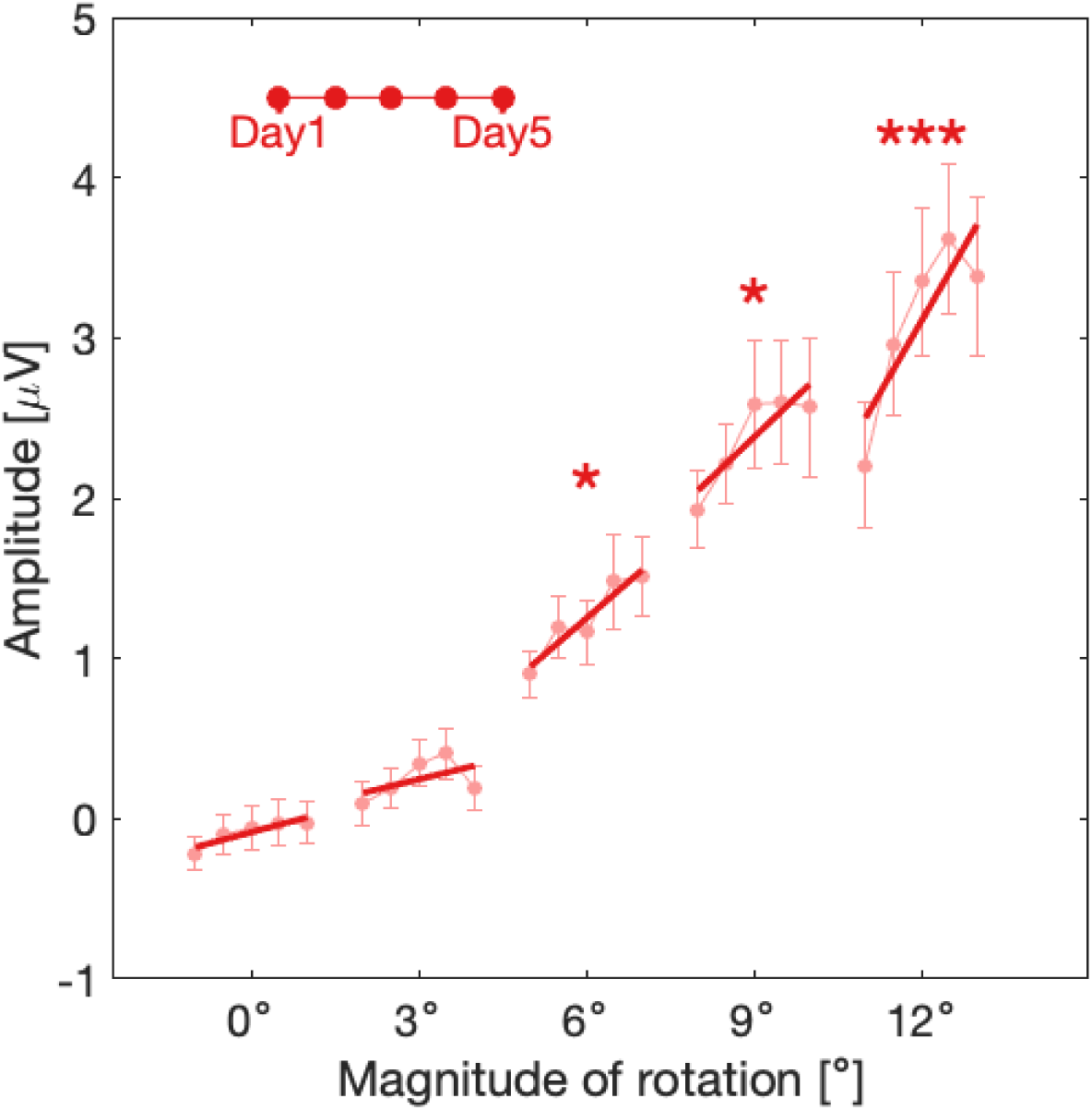
Pe amplitudes across days of perceptual training in the Behavior group (Experiment 1). Pe amplitude at the Cz electrode for each rotation magnitude and training day in the Behavior group (N=16). Error bars represent the mean and standard error across participants. Lines represent the best-fit trend generated using the trust-region algorithm for visualization purposes only and not for statistical testing. Asterisks (*) indicate rotation magnitudes where the fixed effect of training day was significant in the linear mixed-effects models. ∗*P <* 0.05, ∗ ∗ ∗*P <* 0.001.

### BCI-feedback training accelerates perceptual learning of small visuo-motor errors

Figure 5 shows participant’s perceptual accuracy at different rotation magnitudes across training days in the Behavior (Experiment 1) and BCI groups (Experiment 2). In the Behavior group, a two-way rmANOVA with magnitude and days as the main factors revealed a significant main effect of magnitude (*F* (4, 60) = 142.8200, *p_corrected_ <* 0.0001) and days (*F* (4, 60) = 3.5755, *p_corrected_* = 0.0132) and a significant interaction between the two (*F* (16, 240) = 3.4205, *p_corrected_ <* 0.0001), suggesting magnitude-dependent perceptual learning. In the BCI group, a two-way rmANOVA with magnitude and days as main factors revealed the significant main effect of magnitude (*F* (4, 60) = 134.4990, *p_corrected_ <* 0.0001) and days (*F* (4, 60) = 11.5888, *p_corrected_ <* 0.0001). The interaction between the two factors was marginally significant (*F* (16, 240) = 1.5084, *p_corrected_* = 0.0972).

**Figure 5:**
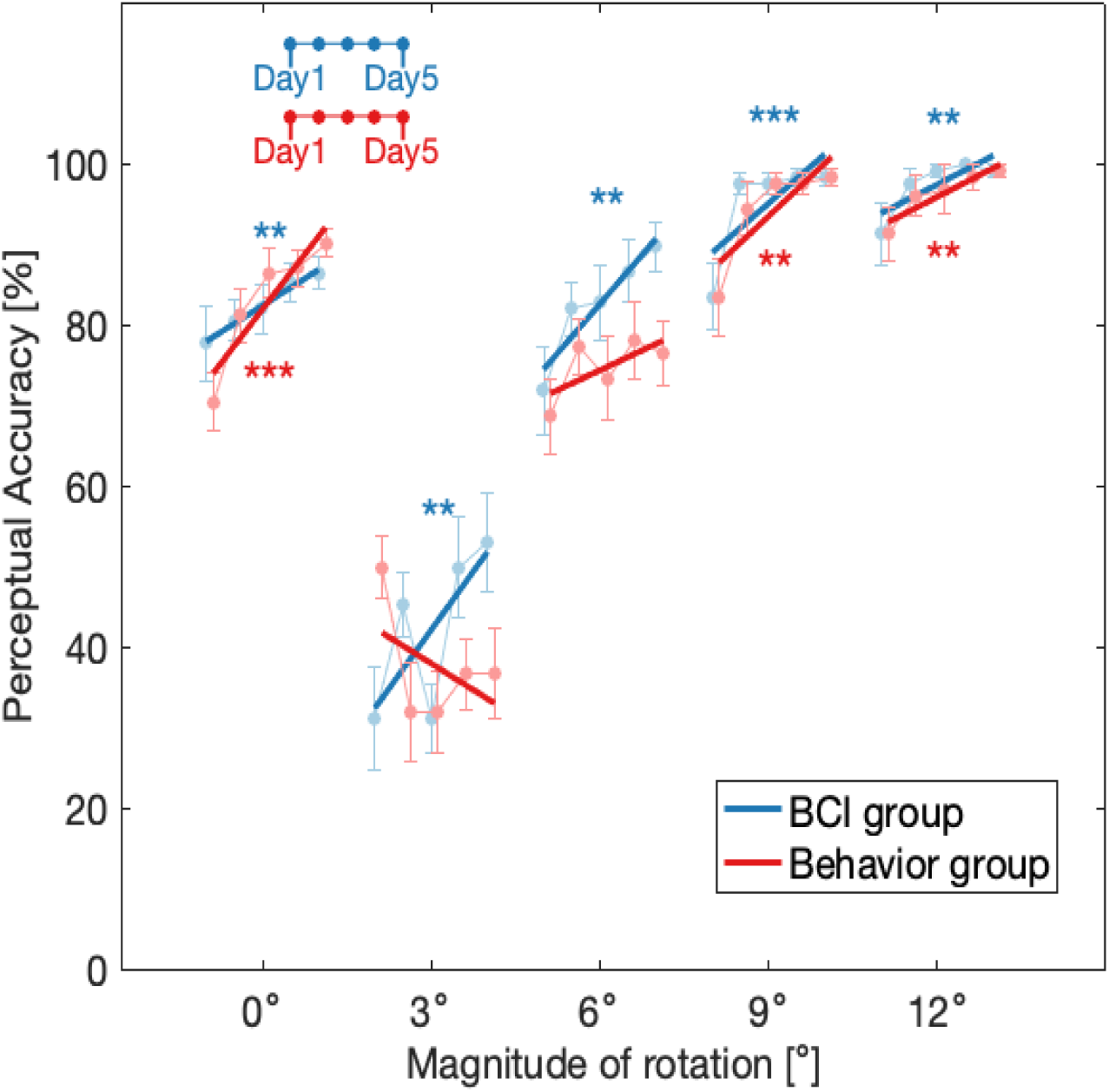
Comparison of perceptual skill performance between groups across training days. Perceptual accuracy at each error magnitudes (0^◦^, 3^◦^, 6^◦^, 9^◦^, 12^◦^) for each each group across the intervention days. The data points show the mean perceptual accuracies across participants, with error bars indicating the standard error. Solid-lines represent the best-fit trend generated using the trust-region algorithm for visualization purposes only and not for statistical testing. Asterisks (*) indicate rotation magnitudes where the fixed effect of training day was significant in the linear mixed-effects models. ∗ ∗ *P <* 0.01 and ∗ ∗ ∗*P <* 0.001.

Mixed-effects modeling of changes in perceptual accuracy relative to Day 1 at 0^◦^, with days as the within-subjects factor and group as the between-subjects factors, was performed to assess training effects across the 5-day intervention. This revealed no significant days × group interaction (*F* (1, 124) = 0.3467, *p_corrected_* = 0.5570). To assess training effects within each group, we next ran separate linear mixed-effects models across the 5 days. In both groups, performance improved significantly across the 5-days perceptual training, with Behavior group showing a larger slope coefficient (Behavior group: *β*(78) = 4.5313 ± 0.6250, *F* (1, 78) = 52.5620, *p_corrected_ <* 0.0001, *R*^2^(78) = 0.6631; BCI group: *β*(78) = 2.2266 ± 0.8062, *F* (1, 78) = 7.6270, *p_corrected_* = 0.0090, *R*^2^(78) = 0.2548). At small visuo-motor error of 3^◦^, the significant main effect of group from mixed-effects modeling did not survive corrections for comparisons (*F* (1, 124) = 4.7814, *p_corrected_* = 0.1836), and the days x group interaction term did not reach significance (*F* (1, 124) = 0.5475, *p_corrected_* = 0.5570). Separate linear mixed-effects models showed that in the Behavior group, performance declined over days (*β*(78) = −2.1875 ± 1.5164, *F* (1, 78) = 2.0810, *p_corrected_* = 0.1702, *R*^2^(96) = 0.0847), while the BCI group exhibited a significant, positive improvement over days (*β*(80) = 4.8437 ± 1.6645, *F* (1, 78) = 8.47, *p_corrected_* = 0.0070, *R*^2^(78) = 0.1415), with perceptual accuracy increasing from below chance (50%) to above chance after 3 days and remaining above chance thereafter. At 6^◦^, the days x group interaction did not reach significance (*F* (1, 124) = 1.4678, *p_corrected_* = 0.5570). Nonetheless, the differing slopes of perceptual accuracy across days indicate a potential divergence in learning trajectories between the groups. In the Behavior group, performance showed an increasing trend over the 5-days, but did not reach significance (*β*(78) = 1.6406 ± 1.2055, *F* (1, 78) = 1.8521, *p_corrected_* = 0.1775, *R*^2^(78) = 0.1956). In contrast, the BCI group demonstrated a significant improvement in accuracy with a larger slope coefficient for days, suggesting a faster rate of learning (*β*(78) = 4.0625 ± 1.1342, *F* (1, 78) = 12.83, *p_corrected_* = 0.0015, *R*^2^(78) = 0.2743). At larger visuo-motor errors at 9^◦^, there was no significant days × group interaction (*F* (1, 124) = 0.8251, *p_corrected_* = 0.5570). Both groups showed significant performance improvements across days, with similar slope coefficient for days (Behavior group: *β*(78) = 3.2813 ± 0.8892, *F* (1, 78) = 13.6160, *p_corrected_* = 0.0014, *R*^2^(78) = 0.1454; BCI group: *β*(78) = 3.0469 ± 0.7285, *F* (1, 78) = 17.4900, *p_corrected_* = 0.0004, *R*^2^(78) = 0.1794). Similarly, at 12^◦^, no significant days × group interaction (*F* (1, 124) = 0.3678, *p_corrected_* = 0.5570) was observed. Both groups improved significantly over the days, with similar slope coefficient for days (Behavior group: *β*(80) = 1.7969 ± 0.6038, *F* (1, 78) = 8.8559, *p_corrected_* = 0.0070, *R*^2^(78) = 0.3874; BCI group: *β*(78) = 1.7969 ± 0.6200, *F* (1, 78) = 8.3985, *p_corrected_* = 0.0070, *R*^2^(78) = 0.0950).

To better understand the start-to-end impact of the interventions on perceptual accuracy in small visuo-motor errors, 3^◦^ and 6^◦^, we built a mixed effects model. The dependent variable was the initial (Day 1) and final perceptual (Day 5) accuracy over the 5-day intervention, with days as the within-subjects factor and group as the between-subjects factor. The mixed effects model revealed a significant days x group interaction for 3^◦^ (*F* (1, 60) = 12.2030, *p_corrected_* = 0.0018), suggesting group-dependent perceptual learning. Within-group comparisons of the Day 1 and Day 5 perceptual performances showed that the Behavior group had a significant decrease in perceptual accuracy, with a larger-than-medium effect size (Day 1 performance: 50 ± 15.138%, Day 5 performance: 36.719 ± 22.578%*, t*(15) = −2.4026*, p_corrected_* = 0.0396*, Cohen*^t^*s d_z_* = −0.69096). Conversely, the BCI group demonstrated a significant increase in perceptual accuracy with a large effect size (Day 1 performance: 31.25 ± 25.82%, Day 5 performance: 53.125 ± 24.367%*, p_corrected_* = 0.0396*, Cohen*^t^*s d_z_* = 0.87138). Furthermore, between-group comparisons of Day 5 perceptual accuracy showed that the BCI group exhibited higher performance with marginal statistical significance (Behavior group: 36.719 ± 22.578%, BCI group: 53.125 ± 24.367%*, t*(30) = 1.976*, p_corrected_* = 0.0767*, Cohen*^t^*s d_z_* = 0.698) (Figure 6A). Unpaired comparisons of improvement in perceptual accuracy (Day 5 minus Day 1) revealed a significantly larger improvement in the BCI group, with a large effect size (Behavior group: −13.281 ± 22.112%, BCI group: 21.875 ± 35.208%, *t*(30) = 3.382*, p_corrected_* = 0.0040*, Cohen*^t^*s d_z_* = 1.196) (Figure 6B). Interestingly, the Behavior group had an initial perceptual advantage at 3^◦^, with a large effect size (Behavior group: 50.0 ± 15.1383%, BCI group: 31.2500 ± 25.8199%, *p_corrected_*= 0.0314*, Cohen*^t^*s d_z_* = 0.8859). Nevertheless, as a result of training, the BCI group not only closed the gap but ultimately surpassed the Behavior group.

**Figure 6:**
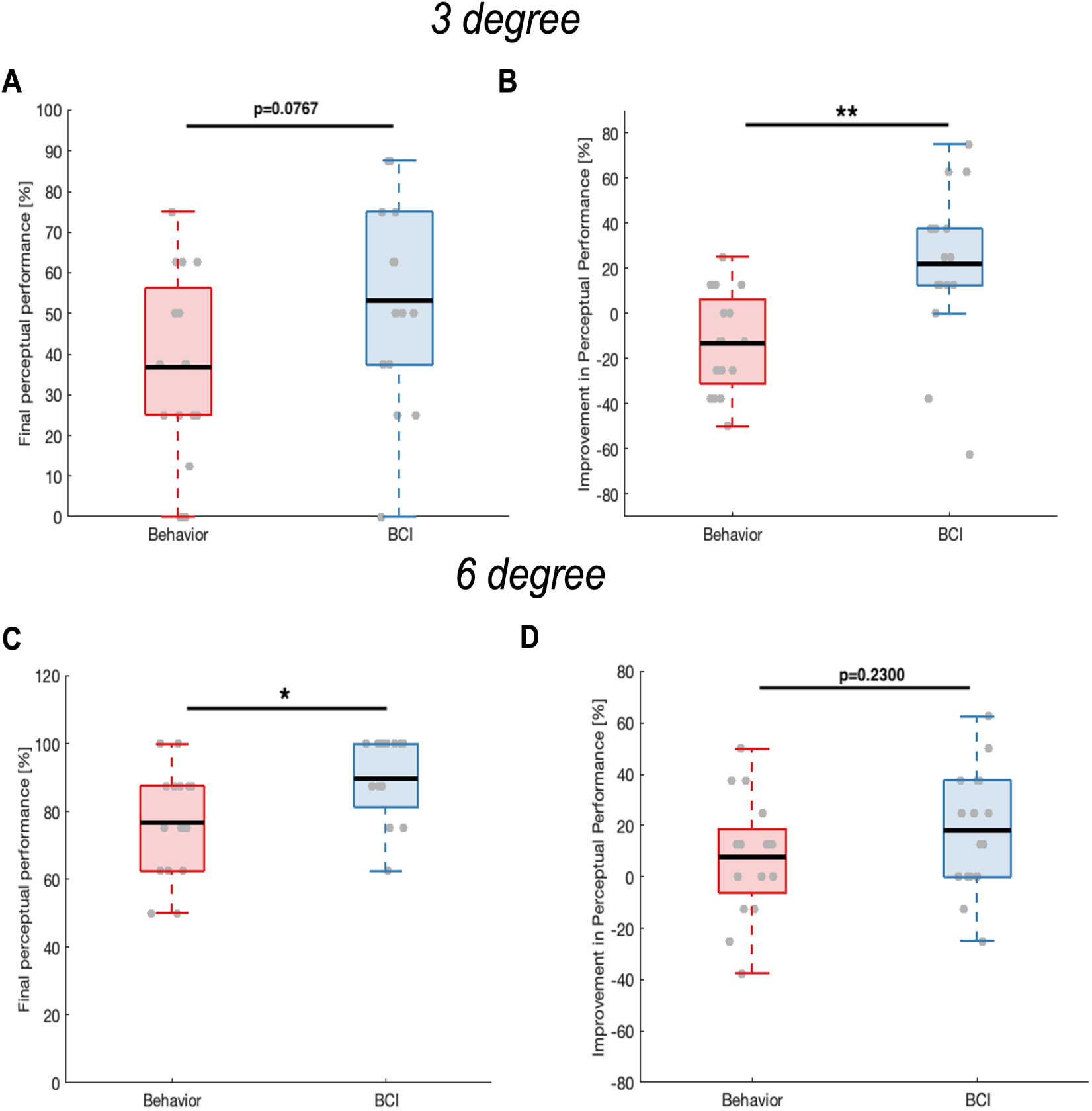
Comparison of perceptual skill performance for small visuo-motor rotations. A: Comparison of perceptual accuracy at 3^◦^ on the final day (Day 5) of training between the Behavior (red, N=16) and BCI (blue, N=16) groups. Boxplots show the distribution of perceptual accuracy across participants, with each data point representing an individual participant. **B**: Comparison of training-induced improvements in perceptual accuracy, defined as the difference between Day 5 and Day 1 accuracy scores, at 3^◦^ between the two groups. Each data point corresponds to one participant. **C**: Comparison of perceptual accuracy at 6^◦^ on the final day of training (Day 5) between the two groups. **D**: Comparison of training-induced improvement in perceptual accuracy at 6^◦^ between the two groups. Between-group comparisons used unpaired tests. ∗*P <* 0.05, ∗ ∗ *P <* 0.001.

For 6^◦^, the days x group interaction did not reach statistical significance (*F* (1, 60) = 1.5991, *p_corrected_* = 0.211). However, the BCI group exhibited a clear trend of perceptual improvement, reflected in a large within-group effect size that was not observed in the Behavior group (Day 1: 71.88 ± 22.13%, Day 5: 89.84 ± 12.26%, *p_corrected_* = 0.0396, Cohen’s *d_z_* = 1.0045). In contrast, within-group analysis in the Behavior group did not show significant improvement in perceptual accuracy from Day 1 to Day 5, with a near-medium effect size (Day 1: 68.75 ± 18.82%, Day 5: 76.56 ± 15.73%, *p_corrected_* = 0.2300, Cohen’s *d_z_* = 0.450). Unpaired comparisons indicated that the perceptual accuracy for 6^◦^ was comparable between the two groups with a small effect size at Day 1 (Behavior: 68.750 ± 18.819%, BCI: 71.875 ± 22.127%, *p_corrected_* = 0.6162*, Cohen*^t^*s d_z_* = 0.152), but it became significantly higher in the BCI group on Day 5 with a large effect size (Behavior: 76.562 ± 15.729%, BCI: 89.844 ± 12.263%, *p_corrected_*= 0.0314*, Cohen*^t^*s d_z_* = 0.942) (Figure 6C). Lastly, an unpaired comparison showed a greater improvement in the BCI group compared to the Behavior group (Figure 6D). The difference was not statistically significant, with a near-medium effect size (Behavior: 7.813 ± 23.218%, BCI: 17.969 ± 23.703%, *t*(30) = 1.2244*, p_corrected_* = 0.2303*, Cohen*^t^*s d_z_* = 0.433).

In addition, we evaluated subject-level rates of perceptual improvement across training days at 3^◦^ and 6^◦^. Each participant’ perceptual abilities were individually fitted using linear regression. At 3^◦^, 14 out of 16 participants (87.5%) in the BCI group showed a positive rate of improvement, in comparison to 7 (43.75%) in the Behavior group. At 6^◦^, 13 in the BCI group (81.25%) showed a positive improvement, in comparison to 10 (62.5%) in the Behavior group. Unpaired comparisons showed significantly higher rates of perceptual improvements in the BCI group than in the Behavior group at 3^◦^ with a large effect size (Behavior: −0.022 ± 0.054, BCI: 0.048 ± 0.076, *p_corrected_*= 0.0030*, Cohen*^t^*s d_z_* = 1.064). For 6^◦^, the BCI group showed higher rates than the Behavior group, with a medium effect size (Behavior: 0.016 ± 0.053, BCI: 0.041 ± 0.051, *p_corrected_*= 0.1970*, Cohen*^t^*s d_z_* = 0.467), although the result did not reach statistical significance.

Lastly, to evaluate the full effect of 5 days of training in the BCI group, we also studied the 6-day trends in perceptual accuracy, which showed consistent trends with the 5-day results. The two-way rmANOVA with magnitude and days as main factors revealed the significant effect of magnitude (*F* (4, 60) = 149.102, *p <* 0.0010) and days (*F* (5, 75) = 12.761, *p <* 0.0010), and no significance in their interactions (*F* (20, 300) = 1.501, *p* = 0.188). Similar to the 5-day results, significant increases in performance were observed across all five rotation magnitudes: 0^◦^: *β*(96) = 2.0759 ± 0.5749, *F* (1, 94) = 13.04, *p_corrected_* = 0.0023, *R*^2^(96) = 0.3212; 3^◦^: *β*(96) = 3.8839 ± 1.2006, *F* (1, 94) = 10.47, *p_corrected_* = 0.0034, *R*^2^(96) = 0.1774; 6^◦^: *β*(96) = 3.5714 ± 0.7944, *F* (1, 94) = 20.21, *p_corrected_* = 0.0005, *R*^2^(96) = 0.3420; 9^◦^: *β*(96) = 2.0982 ± 0.5286, *F* (1, 94) = 15.76, *p_corrected_* = 0.0015, *R*^2^(96) = 0.1410; 12^◦^: *β*(96) = 1.3839 ± 0.4306, *F* (1, 94) = 10.33, *p_corrected_* = 0.0023, *R*^2^(96) = 0.0971).

### Decoding perceptual decisions of visuo-motor errors through BCI

Figure 7 shows the BCI’s online decoding accuracy of the presence or absence of ErrPs at each rotation magnitude. Decoding accuracy for 0^◦^, 6^◦^, 9^◦^, and 12^◦^ consistently remained above chance level (50%) across training days, while performance at 3^◦^ reached chance level by day 2 and stayed above it thereafter.

**Figure 7:**
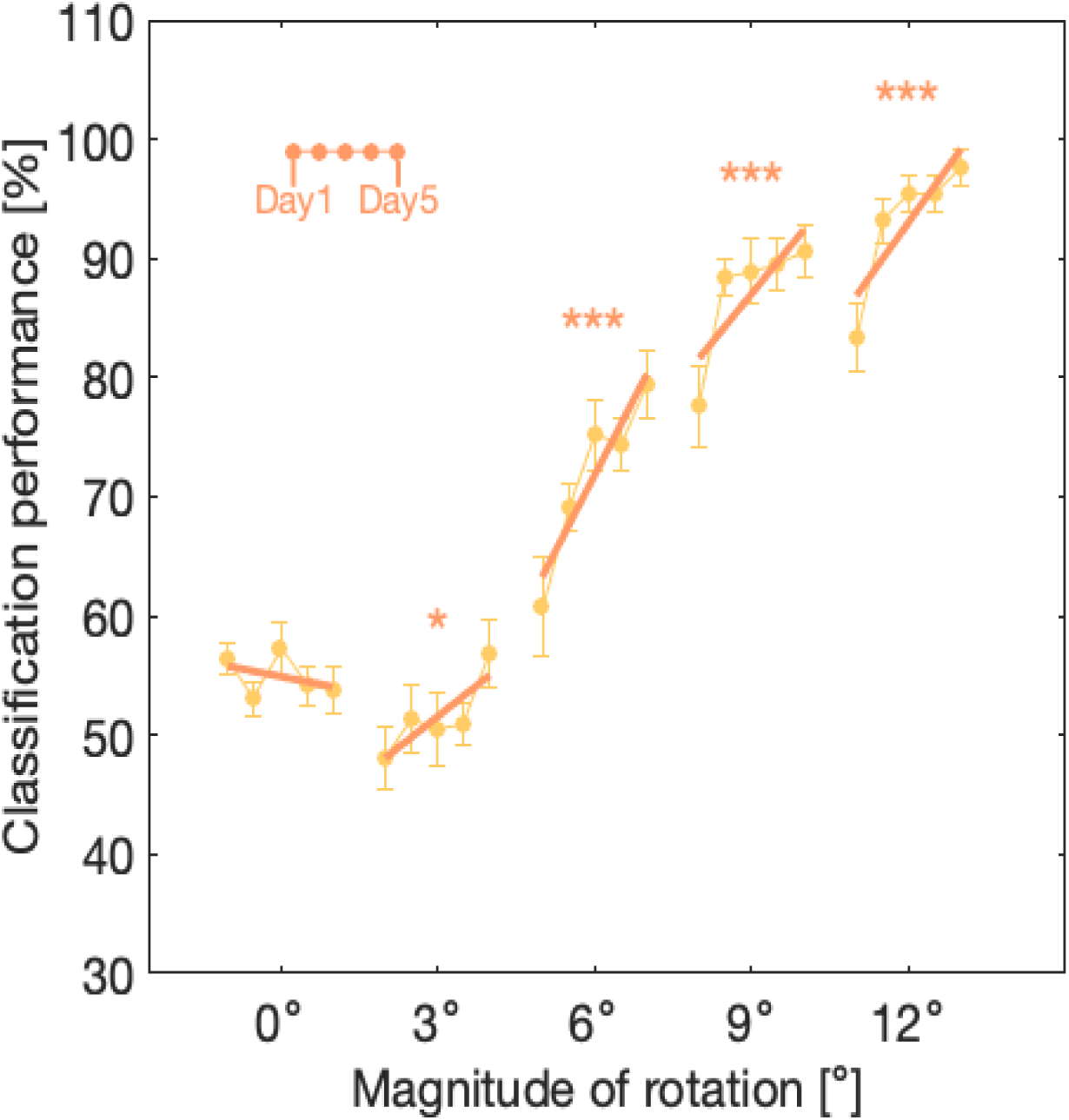
Online decoding accuracy of the presence or absence of ErrPs (Experiment 2). Online decoding accuracy of the presence or absence of ErrPs at each rotation magnitude (0^◦^, 3^◦^, 6^◦^, 9^◦^, 12^◦^) in the BCI group (N=16) across the training days. Data points represent the mean classification performance across participants, with error bars indicating the standard error. Solid-lines represent the best-fit trend generated using the trust-region algorithm for visualization purposes only and not for statistical testing. Asterisks (*) indicate rotation magnitudes where the fixed effect of training day was significant in the linear mixed-effects models. ∗*P <* 0.05 and ∗ ∗ ∗*P <* 0.001.

A two-way rmANOVA with magnitude and days as main factors revealed the significant main effect of magnitude (*F* (4, 60) = 192.3200, *p <* 0.0001) and days (*F* (4, 60) = 15.6000, *p <* 0.0001), and interaction between the two (*F* (16, 240) = 3.4071, *p* = 0.0034). Significant increases in decoding performance were observed over days for 3^◦^, 6^◦^, 9^◦^, and 12^◦^ rotations: 3^◦^: *β*(78) = 1.7379 ± 0.8258, *F* (1, 78) = 4.4289, *p_corrected_* = 0.0483, *R*^2^ = 0.0525; 6^◦^: *β*(78) = 4.2550 ± 0.6946, *F* (1, 78) = 37.5304, *p_corrected_* = 0.0002, *R*^2^ = 0.5630; 9^◦^: (*β*(78) = 0.0270 ± 0.0058, *F* (1, 78) = 21.4471, *p_corrected_* = 0.0002, *R*^2^ = 0.5296; and 12^◦^: *β*(78) = 3.0781±0.4764, *F* (1, 78) = 41.7476, *p_corrected_* = 0.0002, *R*^2^ = 0.5512. In contrast, 0^◦^ showed a non-significant trend in decoding performance (*β*(78) = −0.4320 ± 0.5442, *F* (1, 78) = 0.6329, *p_corrected_* = 0.2000, *R*^2^ = 0.0078).

### BCI feedback during perceptual training enhances the Pe component at small visuo-motor errors

Figure 8A,B illustrate that Pe amplitude was positively correlated with perceptual accuracy for small visual errors. At 3^◦^, the correlation was marginally significant after corrections (Spearman’s *r*(28) = 0.33, *p_corrected_* = 0.0855) while a significant correlation was observed at 6^◦^ (Spearman’s *r*(28) = 0.24, *p_corrected_* = 0.0406).

**Figure 8:**
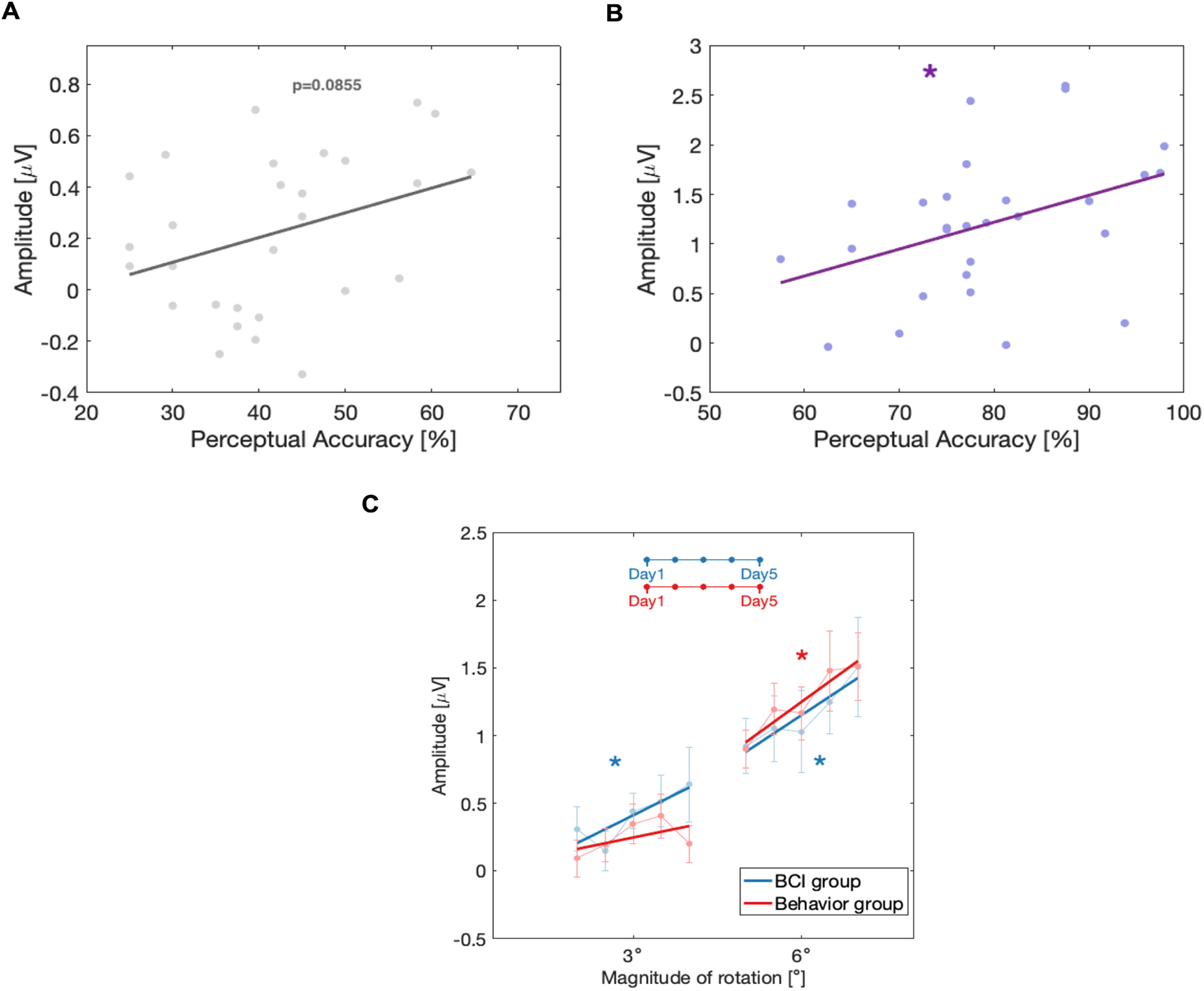
Pe results at Cz. A: Spearman correlation analysis (two-tailed) examining the relationship between Pe amplitudes and perceptual accuracy at 3^◦^ across both groups (N=32). Each data point represents a participant’s average Pe amplitude and perceptual accuracy over the 5 training days. ***B***: Similar analysis for 6^◦^, showing the relationship between Pe amplitudes and perceptual accuracy in both groups. ***C***: Changes in Pe amplitude of ErrPs at 3^◦^ and 6^◦^ for each group across intervention days. Data points represent the mean Pe amplitude at Cz for each intervention day, with error bars indicating the standard error across participants. Solid lines represent the best-fit trend generated using the trust-region algorithm for visualization purposes only and not for statistical testing. Asterisks (*) indicate rotation magnitudes where the fixed effect of training day was significant in the linear mixed-effects models. ∗*P <* 0.05.

We used a mixed-effects model to assess group differences in Pe amplitude changes across days, relative to Day 1, modeling day as a within-subjects factor and group as a between-subjects factor. Our analysis revealed a significant days × group interaction at 3^◦^ that did not survive corrections (*F* (1, 124) = 3.7333, *p_corrected_* = 0.1112), suggesting a potential divergence in Pe amplitude trajectories between groups. To explore within-group changes, we then fit separate linear mixed-effects models across the 5-day period (Figure 8C). In the BCI group, Pe amplitude at 3^◦^ increased significantly across days (*β*(78) = 0.1032 ± 0.0468, *F* (1, 78) = 4.8657, *p_corrected_* = 0.0404, *R*^2^(80) = 0.3524), whereas no significant change was observed in the Behavior group (*β*(78) = 0.0426 ± 0.0387, *F* (1, 78) = 1.2075, *p_corrected_* = 0.2752, *R*^2^(78) = 0.1681). This pattern parallels the group differences in perceptual accuracy shown in Figure 5.

The days × group interaction did not reach significance at 6^◦^ (*F* (1, 124) = 0.1085, *p_corrected_*= 0.7424), indicating consistent trends in Pe amplitudes between the groups. Subsequent linear mixed-effects analysis showed a significant increase in the Pe component over days in both groups (Behavior group: *β*(78) = 0.1508 ± 0.0515, *F* (1, 78) = 8.5578, *p_corrected_* = 0.0156, *R*^2^(78) = 0.4571; BCI group: *β*(78) = 0.1371 ± 0.0502, *F* (1, 78) = 7.4650, *p_corrected_* = 0.0156, *R*^2^(80) = 0.6833).

### Spatial and temporal contributions to ErrP decoding in the BCI group

Since our hypothesis revolves around Pe, the remaining question to ask is which are the main spatial and temporal features that drive BCI feedback —or, in other terms, the spatial and temporal features that are more discriminant in differentiating the presence or absence of an ErrP in the BCI group. In the spatial domain, analyses of the z-scored spatial weights from canonical correlation analysis identified Cz (*Cohen*^t^*s d_z_* = 0.458), CP4 (*Cohen*^t^*s d_z_* = 0.794), and O2 (*Cohen*^t^*s d_z_* = 0.513) as the channels with the largest positive effect sizes among the recorded channels and statistically larger than zero (Figure 9A). However, these significances did not survive multiple comparison corrections. Figure 9B,C confirm physiologically valid grand average ErrP waveforms at CP4 and O2, showing statistically significant differences between error and correct classes in the ERN and Pe regions. Additionally, the contribution of motor areas to ErrP decoding appears minimal, with small or negative effect sizes observed in channels C3 (*Cohen*^t^*s d_z_* = −0.596), C4 (*Cohen*^t^*s d_z_* = 0.116), FC3 (*Cohen*^t^*s d_z_* = −0.396), FC4 (*Cohen*^t^*s d_z_* = −0.194), and CP3 (*Cohen*^t^*s d_z_* = 0.062). These low or negative values suggest that raw spatial weights in these motor channels are close to or below the mean of spatial weights across all recorded channels.

**Figure 9:**
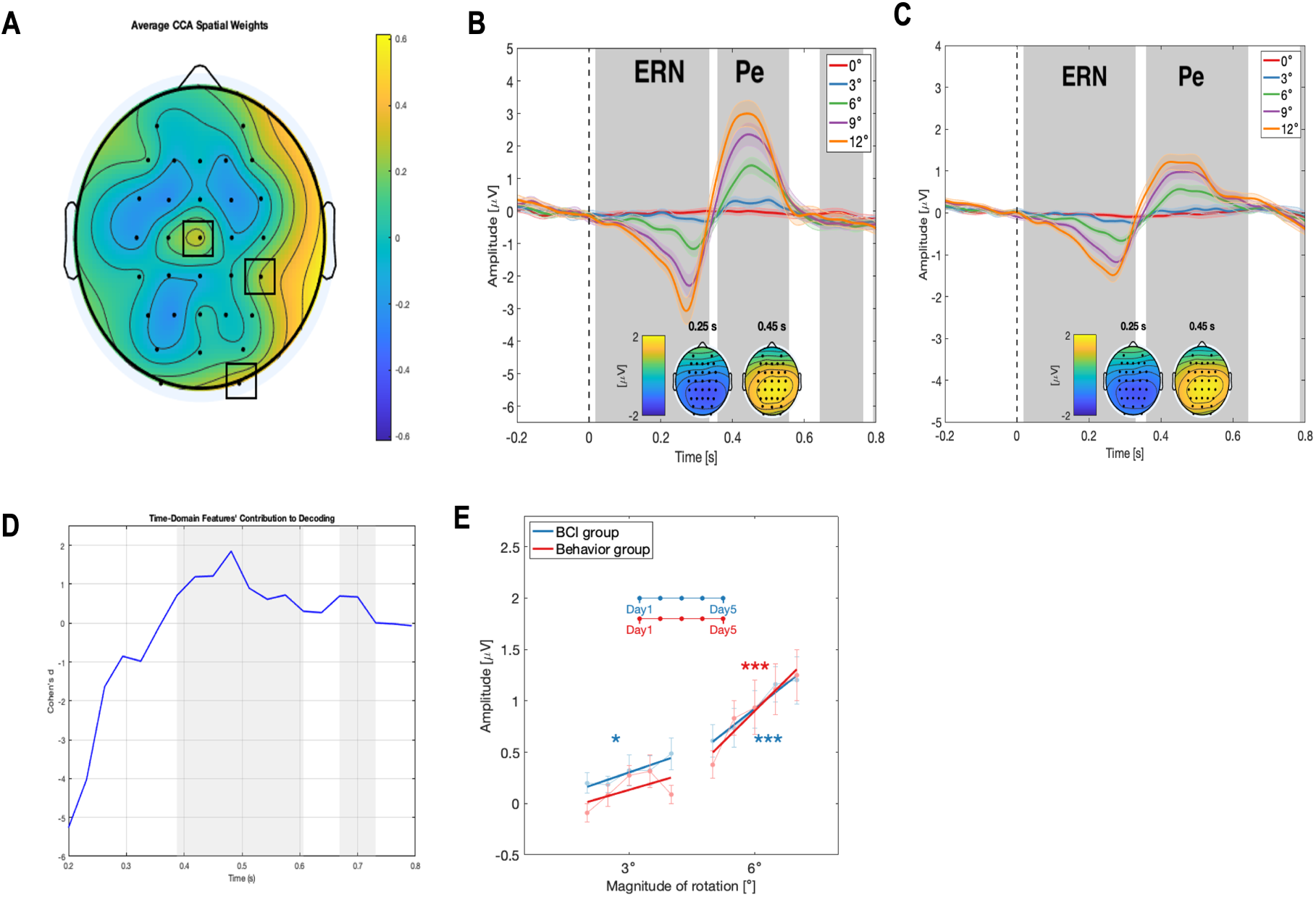
Decoding analyses in the BCI group. ***A***: Topographical plot of the average absolute valued, z-scored CCA, spatial weights across all participants in the BCI group (N=16). Black squares highlight channels with statistically significant spatial weights higher than 0, before multiple corrections: Cz, CP4, O2. ***B*** and ***C***: The time-locked grand average EEG potentials at channel CP4 and O2, respectively, for all participants in the BCI group at different rotation magnitudes (0^◦^, 3^◦^, 6^◦^, 9^◦^, 12^◦^). Time 0 s, marked by horizontal black dashed line, corresponds to the onset of rotation. The shaded areas around the EEG potentials represent the standard error over participants. Gray shaded regions highlight time intervals where significant differences were found between the error and correct class epochs. ***D***: Effect sizes of each absolute valued, z-scored, time-domain feature weight, with the x-axis representing the features’ timestamps. Gray region represents features with weights significantly larger than zero. ***E***: Changes in Pe amplitude at 3^◦^ and 6^◦^ for each group across intervention days at CP4. The data points show the mean Pe amplitudes at day 1-5 of intervention. The error bars show the standard error across participants. Solid lines represent the best-fit trend generated using the trust-region algorithm for visualization purposes only and not for statistical testing. Asterisks (*) indicate rotation magnitudes where the fixed effect of training day was significant in the linear mixed-effects models. ∗*P <* 0.05, ∗ ∗ ∗*P <* 0.001.

In the time domain, a total of nine time-domain features exhibited weights significantly different from zero, as shown in Figure 9D. Of these, seven were located around the Pe region identified in Figure 2. All features in this region demonstrated effect sizes greater than moderate (*Cohen*^t^*s d_z_ >* 0.6), with the highest effect sizes observed in the [418, 481] ms interval, exceeding *Cohen*^t^*s d_z_ >* 1. Interestingly, two features showed significantly positive weights in the N2 region. No frequency-domain analyses were conducted, as PSD features were extracted solely in the [4–8] Hz theta band, which has been shown to characterize ErrPs^27^.

At CP4, mixed-effects model to assess group differences in Pe amplitude changes across days with day as a within-subjects factor and group as a between-subjects factor revealed no significant day × group interaction for 3^◦^ (*F* (1, 124) = 2.2604, *p_corrected_* = 0.2706) or 6^◦^ rotations (*F* (1, 124) = 0.0797, *p_corrected_* = 0.7782). We conducted exploratory linear mixed-effects analyses to examine within-group trends of Pe over days at CP4 given prior evidence suggesting differential Pe evolution at Cz. Figure 9E revealed a significant increase in Pe amplitude at 3^◦^ in the BCI group, while the Behavior group showed marginal significance (Behavior group: *β*(80) = 0.0594 ± 0.0308, *F* (1, 78) = 3.7090, *p_corrected_* = 0.0578, *R*^2^(80) = 0.2110; BCI group: *β*(80) = 0.0701 ± 0.0291, *F* (1, 78) = 5.8157, *p_corrected_* = 0.0243, *R*^2^(80) = 0.4873). At 6^◦^, Pe increased significantly in both groups (Behavior group: *β*(80) = 0.2018 ± 0.0402, *F* (1, 78) = 23.0920, *p_corrected_ <* 0.0001, *R*^2^(80) = 0.6708; BCI group: *β*(80) = 0.1607 ± 0.0393, *F* (1, 78) = 16.7320, *p_corrected_* = 0.0002, *R*^2^(80) = 0.5865).

At O2, no significant day × group interaction was observed at 3^◦^ (*F* (1, 124) = 0.0633, *p_corrected_* = 0.9020) and 6^◦^ (*F* (1, 124) = 0.0152, *p_corrected_* = 0.9020). The grand average for 3^◦^ reveals a nearly flat waveform, indicating a negligible Pe response and making it challenging to estimate Pe at this rotation (Figure 9C). Consequently, linear mixed-effects analyses were not conducted for each group at this channel and magnitude. At 6^◦^, both groups showed significant increases in Pe amplitude (Behavior group: *β*(80) = 0.0465 ± 0.0193, *F* (1, 78) = 5.7092, *p_corrected_* = 0.0193, *R*^2^(80) = 0.5929; BCI group: *β*(80) = 0.0488 ± 0.0199, *F* (1, 78) = 6.0136, *p_corrected_* = 0.0193, *R*^2^(80) = 0.6446).

## Discussion

Neural engineering interventions that target directly the sensory cortices have been used to enhance sensory discrimination in the visual and auditory domains. For instance, a neurofeedback paradigm targeting peak alpha frequency at Pz improved the outcome of perceptual learning in a 3D multiple-object tracking task^34^. Similarly, fMRI-based real-time feedback training enhanced brain activation patterns in early visual areas corresponding to a target orientation stimulus, leading to visual perceptual learning of that pattern ^35^. Other approaches relied on brain or peripheral nerve stimulation. Occipital trancranial alternating current stimulation (tACS) at 10 Hz accelerated perceptual learning in an orientation discrimination task^36^. In the auditory domain, alpha tACS at temporal and central scalp sites improved near-threshold auditory perception relative to sham^37^. Furthermore, vagus nerve stimulation that activated cortically projecting cholinergic axons, improved auditory discrimination performance^38^. An alternative to targeting the sensory cortices is to leverage neural correlates of decision making, a largely unexplored approach to foster perceptual learning. In fact, prior studies have established that changes in activity patterns within the anterior cingulate cortex (ACC), a region involved in decision-making and cognitive control^39, 40^, are closely linked to improvements in perceptual learning outcomes^41^. Other studies have shown that late components of EEG reflecting post-sensory decision-making processes are associated with perceptual learning^42^. In this context, we focus on the perceptions of errors, a key component of adaptive behavior and learning. Critically, ErrPs, which reflect neural processes of error detection and monitoring, originate primarily from the ACC^43–48^. This positions ErrPs as a compelling neural target for modulating perceptual learning through decision-related mechanisms.

Our findings confirmed prior evidence that the Pe magnitude of ErrPs is larger during conscious awareness of errors compared to missed errors^26, 30^ (Figure 3). Building on this, we showed that, as individuals improve their ability to detect errors through a 5-day conventional behavioral perceptual training, their Pe amplitude increases (Figure 4). These establish Pe as a modifiable neural marker of error perception as supported by its positive correlation with perceptual accuracy (Figure 8A,B). We further observed that the 5-day conventional behavioral perceptual training effectively improved the detection of large visuo-motor errors (9^◦^ and 12^◦^), but not of smaller errors (3^◦^ and 6^◦^). In contrast, a 5-day perceptual training with BCI feedback significantly facilitated learning of small errors (3^◦^ and 6^◦^) (Figure 5) and increased their Pe amplitude (Figure 8).

A potential mechanism underlying this enhancement is operant conditioning, a well-established principle in BCI learning^49–52^ and neurofeedback training^53–55^, where participants use feedback on brain activities to voluntarily modulate neural responses. Evidence from other domains has shown that that operant conditioning can modulate evoked potentials, such as motor-evoked potentials (MEPs), which can be increased in both chronic spinal cord injury patients and healthy individuals^56^, pain-related evoked potential components^57^ and slow cortical potentials^58^, though these studies did not use EEG-based BCI feedback. The ErrP feedback combines internal error detection with external reinforcement. This targeted, immediate, and repeated feedback likely activates error-processing pathways during training, potentially strengthening relevant neural circuits and resulting in larger Pe amplitudes over time. Furthermore, we conjecture that, given Pe’s established association with conscious error awareness, the enhanced Pe amplitude subsequently contributed to the improved perceptual performance observed in the BCI group in comparison to the Behavior group. Future studies should further probe the causal role of Pe in perceptual learning using neuromodulation to upregulate and downregulate its amplitude.

It is possible that the feedback timing played an additional role in the BCI group’s superior perceptual learning in comparison to the Behavior group. Feedback timing is a critical factor in feedback-based learning, with immediate feedback enhancing efficiency of learning compared to delayed feedback^59–61^. Prior research suggests that the delay between measurement of a neural signal and its feedback must be kept short to foster activity-dependent neural plasticity^62^. In conventional perceptual training, feedback was delayed as participants must first communicate their perceptual decision^63^. In contrast, the ErrP-BCI feedback provided real-time inferences of participants’ perceptual decisions, eliminating delays. This immediacy likely contributed to the enhanced learning of small errors observed in the BCI group.

Although our hypothesis focused on the physiologically relevance of the Pe component, we used a decoding window [200, 800] ms in the BCI group to extract time-domain features encompassing both the ERN and Pe components. This approach ensured reliable feedback, critical for effective operant conditioning^62, 64^, by leveraging the combined contributions of Pe and ERN to detect ErrPs. Decoding analyses in Figure 9D confirmed that features with the highest contribution to decoding predominantly originated from the Pe region, with some contribution from the N2 region and none from the ERN region. These findings confirm that while feedback incorporated both components, it primarily targeted Pe.

The decoding analyses in Figure 9A revealed that in addition to Cz, CP4 and O2 were also main contributors to the decoding of presence or absence of ErrPs in the BCI group and these channels exhibited physiologically valid ErrP waveforms (Figure 9B,C). These findings suggest that although the Pe component originates in the ACC —typically associated with fronto-central and central regions such as Cz^27^— its scalp expression during visuo-motor tasks can be distributed across multiple areas, including centro-parietal (CP4) and visual (O2) regions. For instance, prior literature has shown parietal ErrPs and theta band activation in error processing during visuomotor movements^5, 65^, supporting the involvement of CP4. Furthermore, CP4 and O2 may reflect the right hemisphere’s specialization in visuospatial processing^66^, particularly the parietal lobe and temporal-occipital cortex^67^. We further speculate that CP4’s involvement due to its role in contralateral motor control and error monitoring during joystick manipulation with the left thumb. Although between group comparisons via mixed-effect modeling did not reach significance, our results demonstrated that Pe amplitude at CP4 and O2 significantly increased during perceptual training of small visuo-motor errors (Figure 9E). Notably, Pe increased significantly for 3^◦^ at CP4 in BCI group (Figure 9E). Accumulating evidence supports the idea that visual perceptual learning is associated with neurophysiological changes in both the primary visual cortex (V1) and higherorder areas of the visual cortex^68–70^. Studies have also reported activity changes in decision-making regions, including the parietal cortex^70, 71^ and ACC^41^, with the parietal cortex implicated in learning visuo-motor errors^72^. These finding align with our results, supporting the idea of neural plasticity within the distributed error-processing networks, like Cz, CP4 and O2.

Our study may have suffered from potential confounding factors, including the possibility that enhanced joystick maneuverability, which could improve participants’ ability to maintain a straight trajectory, might have contributed to the observed differences in perceptual performance over days or between groups. Cursor-reaching duration was used as a metric for joystick maneuverability because participants with better control are more likely to maintain straighter trajectories, resulting in shorter times to reach the target. Straighter trajectories not only minimize duration but might also enhance the ability to perceive perturbations. Conversely, inefficient or curving trajectories, which require more adjustments, increase the duration and might make perception more difficult. Importantly, we found no significant correlation between the cursor-reaching duration and average perceptual accuracy (Spearman correlation, *r*(28) = −0.1218*, p* = 0.5368), suggesting that joystick control did not influence perceptual performance. Mixed effect modeling revealed no day x group interaction (*F* (1, 124) = 1.2073, *p* = 0.2740) or significant main effect of group (*F* (1, 124) = 1.3903, *p* = 0.2406) in the average cursor-reaching duration. Linear mixed effects analyses of cursor-reaching duration across days showed no significant changes over training days for either the Behavior group (*β*(78) = −0.0345 ± 0.0280, *F* (1, 78) = 1.5257, *p_corrected_* = 0.3292, *R*^2^(80) = 0.5819) or the BCI group (*β*(78) = 0.0195 ± 0.0199, *F* (1, 78) = 0.9640, *p_corrected_* = 0.3292, *R*^2^(80) = 0.7642). When averaged across days, the cursor-control durations were not significantly different between groups (Behavior: 2.4517 ± 0.4483 *s*, BCI: 2.4962 ± 0.4497 *s, p* = 0.8951*, Cohen*^t^*s d_z_* = 0.0989). Collectively, these findings suggest that joystick control skills likely didn’t influence perceptual performance nor contributed to observed changes in perceptual performances within or between groups.

Findings from the present study open several avenues for future research. One potential direction is the direct application of the ErrP-BCI intervention to enhance perceptual performance in real-world environments. For example, motorsport drivers, who must rapidly perceive and respond to subtle dynamic changes on tracks under demanding conditions, could benefit from the perceptual enhancements facilitated by these interventions^73^. Beyond benefits to healthy young adults, future research could explore the translational implications of this intervention for addressing sensory perceptual impairments associated with age-related cognitive decline^74, 75^, neuropsychiatric disorders such as bipolar disorder^76^ and schizophrenia^77^. Compared to conventional, behavioral training, an ErrP-BCI intervention offers a more efficient means of accelerating perceptual learning, while also providing a safer alternative to pharmacological approaches^78^. Future studies could also explore whether the 5-day intervention established here leads to long-lasting improvements in perceptual behavior, potentially extending over weeks or months. Alternatively, the dosage, if any, that would enable durable effects. Finally, our demonstration that ErrPs could be modulated through perceptual training and BCI feedback opens new possibilities for cognitive research. ErrP-based interventions could be particularly useful in conditions where ErrPs serve as critical biomarkers, such as adaptive control deficits in schizophrenia^79^ and overactive performance monitoring in obsessive-compulsive disorder^80^. ErrP-BCI approaches might help restore and strengthen cognitive functions impaired by these conditions.

## Methods

### Participants

This study enrolled thirty-two able-bodied, healthy volunteers, divided equally into two groups: the Behavior group (n = 16) and the BCI group (n = 16). The Behavior group consisted of 2 self-reported females and 14 self-reported males, with a mean age of 26 years (SD = 5.0) and 14 right-handed participants. The BCI group included 7 self-reported females and 9 self-reported males, with a mean age of 25 years (SD = 4.5), and all participants were right-handed. All participants reported no history of neurological problems, no current use of psychoactive medication, normal color vision, and normal or corrected-to-normal visual acuity. The experimental protocol was approved by the local ethics commission (2020-03-0073, Austin, Texas, USA). Participants provided written informed consent, adhering to the Declaration of Helsinki, and received compensation for their study participation. The study protocol is published on ClinicalTrials.gov (NCT05311878) and the CONSORT enrollment flow diagram is provided in Figure 10.

**Figure 10:**
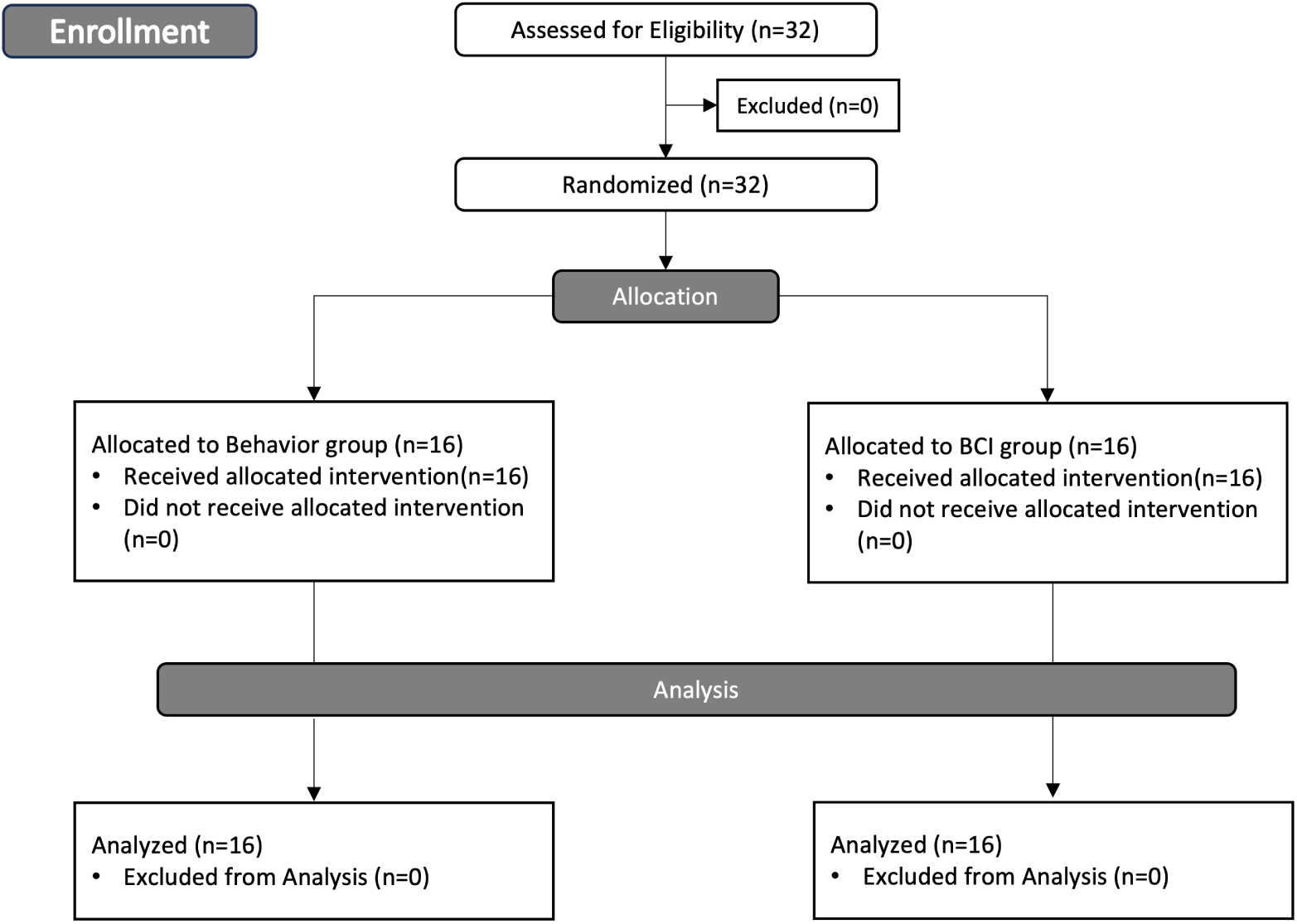
CONSORT flow diagram . Study enrollment diagram. 32 healthy participants were screened and all of them were eligible and agreed to participate. They were randomly assigned to either the Behavior group (n = 16) or the BCI group (n = 16).

### EEG and EOG acquisition

We recorded 32 EEG and 3 electrooculogram (EOG) signals at 512 Hz using an eego system (ANT Neuro, Netherlands). EEG electrodes were located at AF3, AF4, F3, F1, Fz, F2, F4, FC3, FC1, FCz, FC2, FC4, C3, C1, Cz, C2, C4, CP3, CP1, CPz, CP2, CP4, P3, P1, Pz, P2, P4, PO3, POz, PO4, O1 and O2 in 10/20 international coordinates. The ground electrode was placed on the forehead (AFz) and the reference electrode was placed on the inion (Iz). The three EOG electrodes were placed at above the nasion and below the outer canthi of the eyes, and the ground and reference electrodes were placed on the right and left mastoid (M1 and M2), respectively. To reduce signal contamination, participants were asked to avoid excessive eye movements, blinks and motor movements during trials. After visual screening of the EOG signals, participants underwent 90 s of recording at the start of each recording session in which they were asked to perform three different kinds of eye movements for 30 s each: 1) clockwise and counter-clockwise rolling of eyeballs, 2) vertical and horizontal eye movements and 3) repeated eye blinks.

### Experimental procedure

The experimental procedure completed by the two groups is shown in Figure 1C,D. Participants in the Behavior group visited the laboratory for 5 consecutive days, each lasting approximately 90 minutes. Each day began with an EOG calibration run, followed by 10 runs of perceptual training of 32 trials each. Participants in the BCI group attended the laboratory for 6 consecutive days. The first 5 days followed the same structure as the Behavior group with an EOG calibration run and 10 perceptual training runs of 32 trials each. Additionally, each day began with two extra behavioral assessment runs of 32 trials each. These days lasted approximately 110 minutes. The final, 6th day comprised of only two behavioral assessment runs, and lasted approximately 30 minutes. In both groups, EEG data was systematically recorded across all days and runs.

### Experimental setup and task

In this study, participants completed perceptual training in a cursor-reaching task, where errors were induced through visuo-motor rotations—a widely used method known to elicit ErrPs^5, 33, 65^. Participants were seated comfortably in front of a 14-inch display (ThinkPad X1 Carbon, 2560 x 1440 pixels, 60 Hz refresh rate) showing the interface of the cursor-reaching task (Figure 1A,B). The visual scene included a cursor, represented by a blue circle (Diameter: 20 pixels, 0.5 cm), and a target, represented by a red square (80 x 80 pixels, 1 cm x 1 cm). The cursor consistently started from the top-right corner of the display, while the target was positioned at the bottom-left corner, set at a diagonal distance of 9 cm (800 pixels) from the start. Participants used the left joystick of a DualShock4 game-pad (Sony, Japan) to move the cursor from the start to the target. They operated the joystick with minimal left-thumb movements to reduce motor-related EEG artifacts, aiming to minimize these influences on the brain’s perceptual and cognitive processing of errors. The cursor maintained a constant speed of 500 pixels per second (approximately 5.6 cm/s) as long as the joystick was engaged. Participants were instructed to move the cursor in a straight and continuous trajectory, and as swiftly as possible.

In each trial, participants could encounter a potential alteration to the normal joystick-cursor mapping—a visuo-motor error. This change was triggered when the cursor crossed an invisible boundary set at 50% of the diagonal distance between the start and goal locations (Figure 1A,B). The visuo-motor error had one of four possible magnitudes: 3^◦^, 6^◦^, 9^◦^, or 12^◦^. Participants were tasked with detecting this rotation and re-aligning the cursor to its intended straight path.

In both groups, half of the trials in each training day had a visuo-motor rotation, evenly distributed across the four rotation magnitudes and presented in a randomized sequence. They were referred to as the “error” trials. The remaining trials, with a 0^◦^ rotation, served as “correct” trials. In the BCI group’s behavioral assessment runs, the trials were similarly divided, with half designated as “error” trials and the other half as “correct” trials.

### Single-trial procedure

Each trial consisted of the 1) cursor-reaching phase, where participants used the joystick to reach the goal location and the 2) questionnaire and feedback phase. Depending on the experimental group and the nature of the run, this phase could include a questionnaire, feedback, or both.

#### Procedure for the Behavior Group

The sequence of a training trial in the Behavior group is displayed in Figure 1A. Each training trial began with an on-screen text displaying the trial number, after which participants initiated the trial by pressing the X button on the gamepad. Following a randomized delay between 300 and 1000 ms, the cursor and goal locations appeared, marking the start of the cursor-reaching phase. Once the cursor reached the goal location, a 400 ms delay was followed by two on-screen questionnaires. The questionnaires consisted of questions; 1). `Rotation of the mapping?`: whether a mapping rotation occurred in the current trial with the choice of `Yes` or `No`, and 2). `How confident are you?`: participants’ levels of confidence in the former question with responses on a scale from 1 (least confident) to 4 (most confident). Participants selected their answers using the L1, R1, L2, and R2 buttons on the gamepad. Feedback on their perceptual decision was presented 500 ms later, using a green circle (Diameter: 5 cm, 200 pixels) for correct perceptions and a pink X mark (7 cm, 565 pixels per line) for incorrect perceptions. Feedback was presented for 1000ms. Each run ended with a display of the total number of correctly identified trials. Participants were instructed to achieve the highest possible perceptual accuracy in each run.

#### Procedure for the BCI Group

The structure of the trials in the behavioral assessment runs was identical to that of the training trials in the Behavior group, with the exception that participants did not receive feedback on their perceptual performance until the end of the run to prevent learning. Each run ended with a display of the total number of correctly identified trials.

The BCI group’s training trials, displayed in Figure 1B, also ended with a different kind of feedback about perceptual decisions. As shown in Figure 1B, the feedback consisted of: 1). `Rotation`: whether a mapping rotation occurred in the current trial with the choice of `Yes` or `No`, 2). `Error-related potential`: whether the ErrP-BCI detected the presence of an ErrP in the [200, 800] ms period w.r.t the rotation onset, with the decision `Yes` or `No`, and lastly (3) a reward feedback with a green circle (Diameter: 200 pixels, 5 cm) if the decoder output in `Error-related potential` aligned with the rotation mapping in `Rotation`, and a pink X mark otherwise ((7 cm, 565 pixels per line). This feedback was presented for 4,000 ms. Each run ended with a display of a score, calculated as the number of trials in the current run in which ErrP-BCI feedback was aligned with the presence of rotation.

Before the training runs, it was clearly explained that feedback was provided based on the participant’s brain activity. The key instructions were “Please try to achieve the highest possible perception performance in each training run.” and “Whenever you see an incorrect ErrP feedback, please try to assume a mental state/find a strategy where feedback becomes correct in future trials.”

### Decoding the presence/absence of ErrP in the BCI group

During perceptual training, feedback for the BCI group included the output of an online, individually customized BCI, which detected the presence or absence of an ErrP induced by visuo-motor rotations. We used a decoder pipeline similar to prior research^33, 81^. We first applied a 4th order casual Butterworth bandpass filter at 1-10 Hz on EEG and removed EOG artifacts using the autocovariance matrix^82^. EEG signals were then segmented into epochs using window of [200, 800] ms relative to the rotation onset of the cursor. As shown in Figure 2, this window captures the most discriminative time-domain components between the error and correct classes, namely the ERN and Pe components. Epochs with visuo-motor errors (3^◦^, 6^◦^, 9^◦^, or 12^◦^) were labeled as 1 (presence of ErrP), and epochs with 0^◦^ as 0 (absence of ErrP). To enhance the signal-to-noise ratio (SNR) of the EEG signals, we applied a Canonical Correlation Analysis (CCA)-based spatial filter to the epochs, retaining only the first three CCA components for dimensionality reduction^83, 84^. Since ErrPs are characterized by specific temporal signatures (i.e., an increase in signal amplitude in the Pe component relative to baseline) and frequency modulations in the theta band (4-10 Hz)^25, 27^, we extracted two types of features for each CCA component: (1) downsampled signal amplitudes at 32 Hz (yielding 60 temporal features) and (2) power spectral density (PSD) at 4, 6, 8, and 10 Hz frequency bins calculated using Welch’s method (yielding 12 PSD features)^85^. This process produced a feature vector **x** of 72 features per epoch, which was then normalized to the [0, 1] range. The classifier estimated the posterior probability of an ErrP, *p*(*error*|**x**), using a diagonal linear discriminant analysis (LDA), following Equation 1:

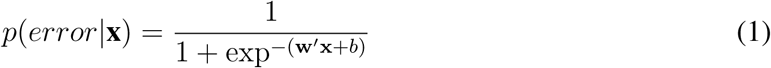

ErrPs were identified if *p*(*error*|**x**) exceeded a decision threshold *τ* ^86^. To ensure robustness and adapt to changes in ErrP characteristics over time, such as those due to learning, the BCI classifier and spatial filters, along with *τ*, were updated after each run using all data from the individual participant.

To optimize *τ* for generalizability across different error magnitudes, we used leave-one-run-out cross-validation, testing *τ* values from 0 to 1 in 0.025 increments. Since the amplitudes of ErrPs encode the magnitudes of error (Figure 2), classifying between error epochs belonging to small rotations (i.e., 3^◦^) and correct epochs (0^◦^) can be challenging. To address this, in cross validation the model built on the training fold was validated on the test fold containing only 0^◦^ correct epochs and 3^◦^ error epochs. The *τ* value that maximized the average True Positive Rate (TPR) and True Negative Rate (TNR) on the average Receiver Operating Characteristic (ROC) curve was selected for real-time use in subsequent runs^86^. TPs were defined as correctly identified ErrPs in error trials and TNs as correctly identified absence of ErrPs in correct trials. Once the optimal *τ* was computed, we constructed the final BCI model using all available participant data for online deployment in the next run.

For the initial perceptual training run of the first day, a personalized decoder was unavailable due to the absence of individual calibration data. Therefore, participants used a generic ErrP decoder, built from all data collected from the 16 participants in the Behavior group. This generic decoder was optimized with leave-one-participant-out cross validation to set the initial *τ* .

### Computation of perceptual accuracy

To evaluate the perceptual accuracy in the BCI group, we calculated it as the percentage of trials correctly perceived at each rotation magnitude in the behavioral assessment runs that commenced each day. Since the Behavior group did not have separate behavioral assessment runs, we matched their initial training trials to the BCI group’s assessment trials by number, rotation magnitude, and day, and evaluated performance on these initial training trials (see Figure 1 C,D). For example, if the BCI group completed eight behavioral assessment trials with a 3^◦^ rotation in day 4, we analyzed the first eight training trials with a 3^◦^ rotation from session 4 in the Behavior group. This enabled direct comparison of perceptual accuracy under identical conditions between the groups.

### Evaluation of joystick handling

To ensure that joystick handling was consistent across training days and between groups, we calculated the average cursor trajectory duration for each participant in each day. This duration was measured from the appearance of the start and goal locations until the cursor reached the goal.

### Error-positivity (Pe) analysis

To analyze the Pe component of the ErrPs, we first applied a non-causal, 4th order bandpass Butterworth filter with cutoff frequencies of 1-10 Hz to extract the theta band activity of the EEG signals. EOG artifacts were then regressed out of the EEG signals using the autocovariance matrix of both the EEG and EOG signals, as described by Schlögl et al.^82^. Next, we segmented the pre-processed EEG signals into epochs using the time window of [-200 1000] ms relative to the onset of rotation. An epoch with any sample exceeding 150 *µV* in any of the recorded channels was considered anomalous and removed. This screening process removed 0.4% of epochs in the Behavior group and 1.4% of epochs in the BCI group. To identify the Pe region, we compared the grand average waveforms of the channel of interest for correct trials (0^◦^) and error trials (3^◦^, 6^◦^, 9^◦^, or 12^◦^). The Pe component was defined as the time window where significant differences between these grand average waveforms were observed. As shown in Figure 2, the Pe component was observed as a positive deflection within the time window of ∼[340, 520] ms relative to the onset. The amplitude of the Pe component was quantified by measuring the mean voltage within this time frame. EEG data processing and analyses were carried out in Matlab (MathWorks, Natick, MA).

### Analyses of spatial and temporal contributions to ErrP decoding in the BCI group

To evaluate the contributions of individual channels to the ErrP decoder in the BCI group, we extracted spatial weights from the CCA, which was used as the spatial filtering method in the decoding pipeline. These spatial weights were absolute valued, z-scored and averaged across components, runs, and days to produce a single weight for each channel per participant. We averaged across runs because the decoder was updated after each run, resulting in a new set of CCA weights.

To analyze the contributions of temporal features to the ErrP decoding, we extracted the LDA weights for time domain features used in each run. These weights were similarly absolute valued, z-scored across CCA components, runs and days, yielding a single weight for each time-domain feature per participant.

### Statistical analysis

All statistical tests were conducted on data that satisfied their assumptions. We evaluated data normality using the Lilliefors test, with a p-value threshold of 0.05 to indicate normal distribution^87^.

To examine the impact of behavioral response (successfully perceived vs missed) on the Pe amplitude at 3^◦^ and 6^◦^ in the Behavior group, we used a two-way repeated-measures ANOVA (rmANOVA), with behavioral response (Perceived vs Missed) and rotation magnitudes (3^◦^, 6^◦^) as within-subject factors. The dependent variable was the average Pe amplitude for each response type and magnitude. This was followed by a paired two-tailed t-tests to compare Pe amplitudes across different perceptual conditions. For non-normally distributed data, Wilcoxon signed-rank tests were used.m

Changes in Pe amplitude across days in the Behavior group were first assessed using a twoway repeated-measures ANOVA with day and rotation magnitude as within-subject factors, reporting F-statistics and p-values. The dependent variable was the average Pe amplitude for each day and magnitude. To model day-wise changes in Pe amplitude at each rotation magnitude, we used linear mixed-effects (LME) models with day as a fixed effect and participant as a random intercept to account for individual variability. Similarly, LME models were used to assess changes in perceptual accuracy and online classification accuracy across days, using the same model structure. For all LME analyses, results were reported using the fixed-effect coefficient and its standard error, the F-statistic, corresponding p-value, and *R*^2^ as a measure of model fit.

We used mixed-effects models to examine the main effects of group and the interaction between day and group on perceptual performance and Pe amplitude, with participants modeled as random intercepts. The dependent variable was defined as the change in each metric relative to its value on Day 1.

Group differences in perceptual accuracy, perceptual improvement, and improvement rates at specific rotation magnitudes were conducted using unpaired two-tailed t-tests. For non-normally distributed data, Mann–Whitney U tests were used. Improvement rates were calculated as the slopes of linear models fitted to individual subject’s data across days. For all paired and unpaired tests, effect sizes were reported using Cohen’s *d_z_*, and t-statistics were provided when applicable to indicate the magnitude of observed effects. Means and standard deviations were also reported to describe central tendency and variability.

Two-tailed Spearman correlations were used to assess relationships between Pe amplitude and perceptual performance, with each data point representing a participant’s average Pe amplitude and accuracy at a given rotation magnitude across training days. Similarly, correlations between perceptual performance and cursor-reaching duration were computed using participant-level averages across all training days and rotation magnitudes. Similar correlations were also computed between Pe amplitude and rotation magnitude. To minimize outlier effects, univariate and bivariate outliers were identified and excluded using Carling’s modified boxplot rule^88, 89^.

To analyze spatial contributions to ErrP decoding in the BCI group, averaged spatial weights were compared to zero using one-tailed paired tests, with corresponding effect sizes computed. Temporal analyses used the same approach, testing LDA feature weights against zero and reporting effect sizes.

To extract Pe components at channels Cz, CP4, and O2, we computed grand-average waveforms for error and correct trials for each of the participants from both groups. Paired comparisons were then performed at each time point within a [0, 1000] ms window, using the participant-level grand averages. Time points showing significant differences after correction were identified as Pe component regions.

All statistical analyses were conducted at the participant level. All analyses were subject to corrections for multiple comparisons using the Benjamini-Hochberg method.

## Author Contributions

D.L., F.I., and J.d.R.M. conceived and designed the experimental protocols. D.L. and F.I. implemented the experimental protocol. D.L., F.I., and M.Z. were responsible for data acquisition. D.L. and F.I. performed the analyses. D.L., F.I., and J.d.R.M. interpreted results. D.L. and F.I. prepared the draft manuscript. All authors reviewed and approved the final version of the manuscript.

